# Nucleotide-Driven Triple-State Remodeling of the AAA-ATPase Channel in the Activated Human 26S Proteasome

**DOI:** 10.1101/132613

**Authors:** Yanan Zhu, Wei Li Wang, Daqi Yu, Qi Ouyang, Ying Lu, Youdong Mao

## Abstract

The proteasome is a sophisticated ATP-dependent molecular machine responsible for protein degradation in all eukaryotic cells. It remains elusive how conformational changes of the AAA-ATPase unfoldase in the regulatory particle (RP) control the gating of substrate-translocation channel to the proteolytic chamber of the core particle (CP). Here we report three alternative states of the ATP-γS-bound human proteasome, in which the CP gate is asymmetrically open, visualized by cryo-EM at near-atomic resolutions. Only four nucleotides are stably bound to the AAA-ATPase ring in the open-gate states. Concerted nucleotide exchange gives rise to a back-and-forth wobbling motion of the AAA-ATPase channel, coincident with remarkable transitions of their pore loops between the spiral staircase and saddle-shaped circle topologies. Gate opening in the CP is thus controlled with nucleotide-driven remodeling of the AAA-ATPase unfoldase. These findings demonstrate an elegant mechanism of allosteric coordination among sub-machines within the holoenzyme that is crucial for substrate translocation.

## INTRODUCTION

The ubiquitin-proteasome system (UPS) participates in numerous important biological processes, such as regulation of gene expression, cell division, innate and adaptive immunity, and the response to proteotoxic stress (Finley, 2009; Finley et al., 2016; Kish-Trier and Hill, 2013; Tomko and Hochstrasser, 2013). A set of ubiquitylation pathways conjugate ubiquitin chains on target proteins, which are recognized and degraded by the 2.5-MDa 26S proteasome holoenzyme. The proteasome is composed of a 28-subunit barrel-shaped core particle (CP) and two 19-subunit regulatory particles (RP) (Beck et al., 2012; da Fonseca et al., 2012; Lander et al., 2012; Lasker et al., 2012) capped at both sides of the CP. The catalytic chamber within the CP contains three proteolytically active threonine residues. Substrate entry into this chamber for degradation is controlled through the axial channel in the center of a heptameric α-ring, also called the CP gate, positioned on each side of the CP. Specific interactions of the α-ring with the RP or other activators may result in the opening the CP gate (Chen et al., 2016; Groll et al., 1997; Harshbarger et al., 2015; Lowe et al., 1995; Unno et al., 2002; Wehmer et al., 2017). The RP consists of two subcomplexes known as the lid and the base (Glickman et al., 1998). Recognition of a ubiquitylated substrate is mediated principally by ubiquitin receptors Rpn10 and Rpn13, the base subunits within the holoenzyme (Finley, 2009). The globular domains of a substrate are mechanically unfolded by a ring-like heterohexameric subcomplex in the base consisting of six distinct subunits, Rpt1-6, which belong to the ATPases-associated-with-diverse-cellular-activities (AAA) family. To allow efficient substrate translocation, conjugated ubiquitins are removed by a metalloprotease subunit Rpn11 in the lid.

Although major advances have been made in the last three decades in understanding the proteasome architecture (Beck et al., 2012; Boehringer et al., 2012; da Fonseca et al., 2012; Dambacher et al., 2016; He et al., 2012; Kish-Trier and Hill, 2013; Lander et al., 2012; Lasker et al., 2012; Lowe et al., 1995; Luan et al., 2016; Pathare et al., 2012; Pathare et al., 2014; Riedinger et al., 2010; Tomko and Hochstrasser, 2013; Wang et al., 2005; Worden et al., 2014; Zhang et al., 2009a; Zhang et al., 2009b; Zhang et al., 2009c), only recently was the complete, intact proteasome in a resting state resolved to near-atomic resolutions, at which a reliable Cα-backbone can be traced with partial assignment of amino acids (Chen et al., 2016; Ding et al., 2017; Huang et al., 2016; Schweitzer et al., 2016; Wehmer et al., 2017). Two other conformational states of the yeast proteasome were also resolved to similar resolutions lately (Ding et al., 2017; Wehmer et al., 2017). However, all these high-resolution structures revealed the CP gate in a closed conformation incompatible with substrate degradation. We recently discovered that the human proteasome can assume four major conformations in a common solution condition with saturated ATP, designated the ground (S_A_), the commitment (S_B_), the gate-priming (S_C_) and the open-gate (S_D_) states (Chen et al., 2016). Only the S_A_ state was resolved to high resolution, whereas the resolutions of the other states remain relatively low.

Except for the S_D_ state, all other states were found to be closed in their CP gates. In addition to earlier studies by cryo-EM revealing that the yeast proteasome may assume three conformational states (s1, s2, and s3) (Beck et al., 2012; da Fonseca et al., 2012; Lander et al., 2012; Lasker et al., 2012; Luan et al., 2016; Matyskiela et al., 2013; Sledz et al., 2013; Unverdorben et al., 2014), the existence of the fourth state s4, analogous to the human S_D_ state, was recently discovered in the yeast proteasome at 7.7-Å resolution (Wehmer et al., 2017), which shared similar structural features as those found in the human proteasome S_D_ state, such as an open gate in the CP. However, the insufficient resolution of the S_D_ or s4 state significantly limit our understanding of the proteasome activation as well as the structural mechanism of substrate degradation (Chen et al., 2016; Wehmer et al., 2017).

Here, we report near-atomic resolution cryo-electron microscopy (cryo-EM) structures of the activated human proteasome in complex with ATP-γS in three alternative conformational states (designated S_D1_, S_D2_ and S_D3_), in which the CP gates are asymmetrically open. The three structures mainly differ in their AAA-ATPase ring, in which the nucleotide exchange is found to drive a wobbling motion in the base. This allows us to visualize remarkable architectural transitions of the pore loops along the substrate-translocation pathway between the spiral staircase and saddle-shaped circle topologies. We also report the ATP-γS-bound human proteasome in the S_A_, S_B_ and S_C_ states, in which the CP gates are closed under the same physiological condition. Taken together, these findings demonstrate an elegant mechanism of allosteric coordination among sub-machines within the holoenzyme that is crucial for substrate translocation.

## RESULTS

### Cryo-EM Structure Determination of the Gate-Opened Proteasome Holoenzyme

We collected cryo-EM data on the human proteasome holoenzyme, supplied with 1 mM ATP-γS, using a Gatan K2 Summit direct electron detector mounted on a 200-kV cryogenic electron microscope Tecnai Arctica (Figure 1A, and Figure S1). Using a deep-learning based method, we automatically extracted 502,384 single-particle images from 8,463 drift-corrected micrographs (Zhu et al., 2016). After unsupervised 2D classification (Figure 2B), 395,941 single-particle images were verified and chosen for further analysis (Wu et al., 2016). The single-particle images of doubly-capped proteasomes were then converted to single-particle images of RP-CP subcomplex by a 3D soft mask excluding one of the two RPs in image alignment and classification. Ultimately, through exhaustive unsupervised focused 3D classification (Scheres et al., 2007), we obtained six conformations that were separately refined to their best resolutions (Figure S2). Three of them correspond to the previously reported S_A_, S_B_ and S_C_ states, which were refined to nominal resolutions of 3.6, 7.0 and 5.8 Å, respectively (Figure 1C, Figures S1, and S3). The other three conformations, S_D1_, S_D2_ and S_D3_, all featuring an open CP gate, were refined to nominal resolutions of 4.2, 4.3 and 4.9 Å, respectively (Figure 1D-F, Figures S1 and S3).

**Figure 1.**
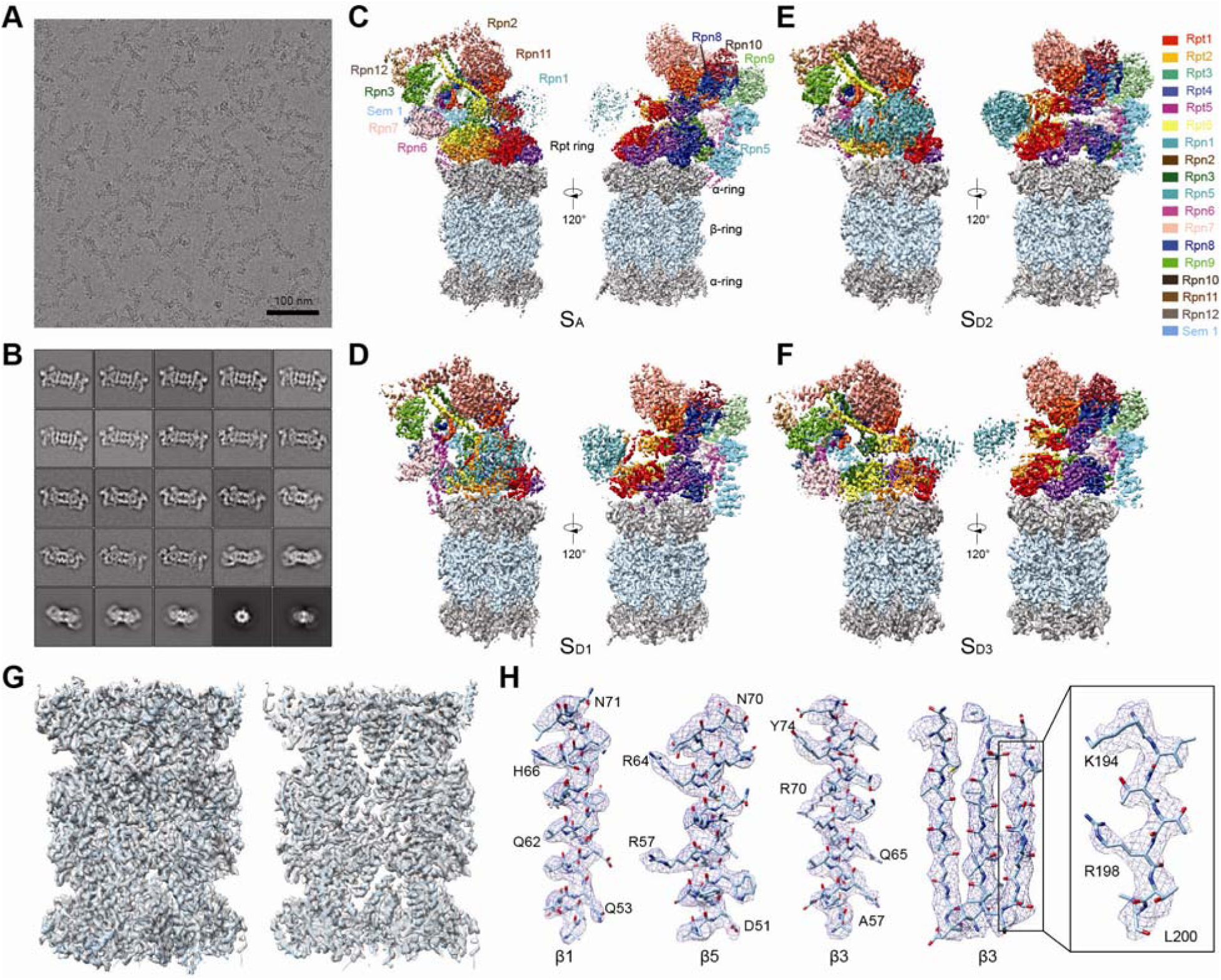
Cryo-EM structure determination of the ATPγS-bound human proteasome. A typical cryo-EM micrograph of the ATP-γS-bound human proteasome imaged with a Tecnai Arctica and Gatan K2 Summit direct detector camera. Typical reference-free 2D class averages computed by the ROME software. (C-F) Two different views of the cryo-EM density maps colored according to its subunits in the S_A_ (panel C), S_D1_ (panel D), S_D2_ (panel E), S_D3_ (panel F) states. The 3.5-Å cryo-EM density of the CP in the combined S_D_ state is superposed with its atomic model. Left, a lateral perspective. Right, a central slice. Typical high-resolution densities of the secondary structures in the cryo-EM structure of the CP in the open-gate S_D_ state.

**Figure 2.**
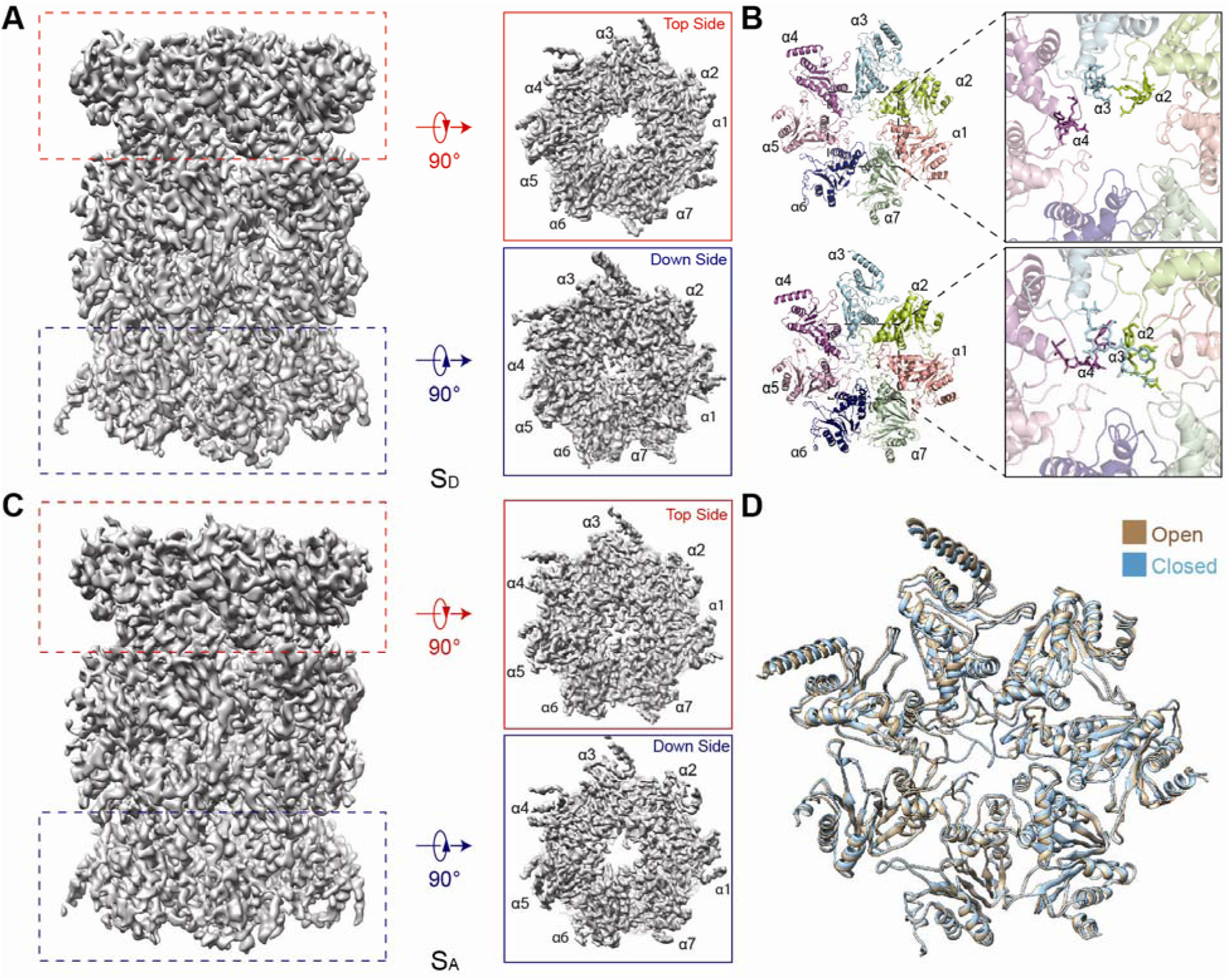
Asymmetric opening of the CP gates in the S_A_ and S_D_ states. The side view (left), top view (upper right) and bottom view (lower right) of the CP density map in the combined S_D_ state. The atomic models of two different α-rings in cartoon representation from the perspective of the AAA-ATPase or the RP-CP interface in S_D_ (left). Close-up view of the central part of the α-rings, showing that the CP gate is open on the top (upper right) and closed on the bottom (lower right). The side view (left), top view (upper right) and bottom view (lower right) of the CP density map in the S_A_ state. Superimposed atomic models of two different α-rings in cartoon representation in the S_D_ state and colored light orange (closed) and light blue (open).

The structures of CP in the S_D1_, S_D2_ and S_D3_ states are virtually identical up to their measured resolution and reproduce the open-gate CP in the 8-Å structure of the S_D_ state reported recently (Chen et al., 2016). Based on this observation, we combined the datasets of S_D1_, S_D2_ and S_D3_ and performed a high-resolution refinement focusing on the CP component, by applying a local mask that retains the CP structure. This allowed us to improve the open-gate CP structure in the S_D1-3_ states to an average resolution of 3.5 Å (Figure 1G, H, Figures S1 and S3). Based on the atomic model of the S_A_ state, we built and refined three atomic models for S_D1_, S_D2_ and S_D3_ (Figures S3, S4). Consistent with the S_A_ structure, the ubiquitin receptor Rpn13 and the ubiquitin-interacting motif (UIM) of the ubiquitin receptor Rpn10 are missing in the cryo-EM densities in all conformational states (Wang et al., 2005; Zhang et al., 2009c).

### Conformational States of the ATP-γS-Bound Human Proteasome

The S_A_ and S_C_ states of the ATP-γS-bound human proteasome are nearly identical to the previously reported S_A_ and S_C_ states of the ATP-bound human proteasome (Figure S5A, C). The S_B_ state also closely resembles the previously reported S_B_ state, but shows a rather small rotation of RP against the interface between Rpn2 and the OB ring (Figure S5B). Consistent with our previous report, the S_A_, S_B_ and S_C_ states of the ATP-γS-bound human proteasome all show a closed CP gate (Chen et al., 2016).

The S_D1_, S_D2_ and S_D3_ states all exhibit defined differences in AAA-ATPase as compared to the previously reported S_D_ state (Chen et al., 2016) (Figure S5D-F). When the three datasets were combined to produce a merged cryo-EM reconstruction, the resulting map closely reproduced the previously reported S_D_ conformation (Chen et al., 2016) (Figure S5G,H). In contrast to the improved resolution of the CP in the combined S_D_ map, the resolution of the AAA-ATPase module is limited to 5-6 Å in the combined S_D_ map, and is lower than those achieved in the separately refined S_D1_, S_D2_ and S_D3_ maps. Moreover, the AAA-ATPase module in the combined reconstruction also exhibits reduced densities as opposed to the lid subcomplex, indicating greater structural heterogeneity in the AAA-ATPase than in other components. Separation of the three datasets, albeit reducing the particle numbers significantly, instead improved the AAA-ATPase resolution. These observations indicate that the previously reported S_D_ state is still heterogeneous, and is most likely an average of multiple conformations (Chen et al., 2016), which are now captured in the S_D1_, S_D2_ and S_D3_ states.

The probability of observing an open gate in the CP was dramatically enhanced in the presence of ATP-γS. The S_A_, S_B_, S_C_ and S_D_ states in the ATP-bound human proteasome were represented in the particle populations at 76.1%, 10.2%, 5.8% and 7.9%, respectively (Chen et al., 2016). In contrast, the states of S_A_, S_B_, S_C_, S_D1_, S_D2_ and S_D3_ are represented in the particle populations at 51.8%, 3.5%, 5.3%, 14.9%, 17.0% and 7.5%, respectively. Thus, the probability for a human proteasome to adopt a S_D_-like state in the presence of ATP-γS is about five times that in the presence of ATP. Binding of the slowly hydrolysable nucleotide analog ATP-γS, instead of ATP, apparently changed the conformational equilibrium among the coexisting states in favor of the S_D_-like states.

### Asymmetric Gate Opening in the CP

The CP structure in the resting S_A_ state of ATP-bound human proteasome exhibits a *C2* symmetry. However, this symmetry is broken in the ATP-γS-bound human proteasome. In the CP of the three S_D_ states, we observed asymmetric gate opening (Figure 2A). While the α-ring in contact with the RP in the S_D_ states is open in the center, the other α-ring in the opposite side of CP remains mostly closed at the same level of the density contour (Figure 2A). Interestingly, in the S_A_ map, where the α-ring in contact with the RP in the S_A_ state is closed, the opposite side of CP is instead mostly open (Figure 2C). Thus, in most ATP-γS-bound proteasome holoenzyme, the CP gate opening does not take place in both α-rings of each holoenzyme simultaneously. Instead, only one of the two gates in each holoenzyme is open at a time on most particles.

The CP gate is principally controlled by the N-terminal tails of the α2 and α4 subunits; and the N-terminal tail of α3-subunit behaves as a lynchpin of the gate, stabilizing the closed state of the CP gate (Groll et al., 2000). Reconfiguration of the α3 tail is controlled by Rpt2 (Tian et al., 2011). The reorientation of these tails constitutes gating and controls substrate entry into the CP. Previous studies suggested that the gate opening in archaeal CP is coincident with a prominent rotation in the α-subunits (Yu et al., 2010). On the contrary, although the N-terminal tails of α2, α3 and α4 are rotated over a large angle to roughly align along the heptameric axis to open the CP gate (Figure 2B), the helical elements connected to the gate-blocking tails in the α-subunits are nearly identical to those in the closed CP (Figure 2D). This observation is consistent with the crystal structures of yeast 20S proteasome in complex with 11S regulators (Stadtmueller et al., 2010; Whitby et al., 2000) as well as the s4 state of the yeast 26S proteasome (Wehmer et al., 2017).

### Conformational Fluctuations of the RP in the Open-Gate States

The RP in the three open-gate states rotates and translates above the CP, with its subunits undergoing differential movements (Figure 3). Relative to the S_D1_ state, the lid subcomplex in S_D2_ is overall translated for 1-2 nm toward the side of Rpn3 and Rpn12 (Figure 3A, B). The hexameric AAA-ATPase ring travels ∼1 nm above the heptameric α-ring in the direction along the lid translation (Figure 3D-I), whereas Rpn1 translated in the opposite direction for about 1 nm. The lid subcomplex further rotates ∼5° clockwise in S_D3_ relative to S_D2_ (Figure 3B, C). By contrast, Rpn1 rotates ∼30° counterclockwise in S_D3_ relative to S_D2_. The AAA-ATPase ring overall moves ∼1 nm backward in the S_D3_-to-S_D2_ transition as opposed to the S_D2_-to-S_D1_ transition. Thus, the AAA-ATPase ring displays a lateral, back-and-forth vibration above the α-ring as the holoenzyme sampling the three open-gate states, while the OB ring tilts up-and-down with a rocking-like vibration. Meanwhile, the lid subcomplex rotates along an asymmetric circular trajectory with a small swinging angle.

**Figure 3.**
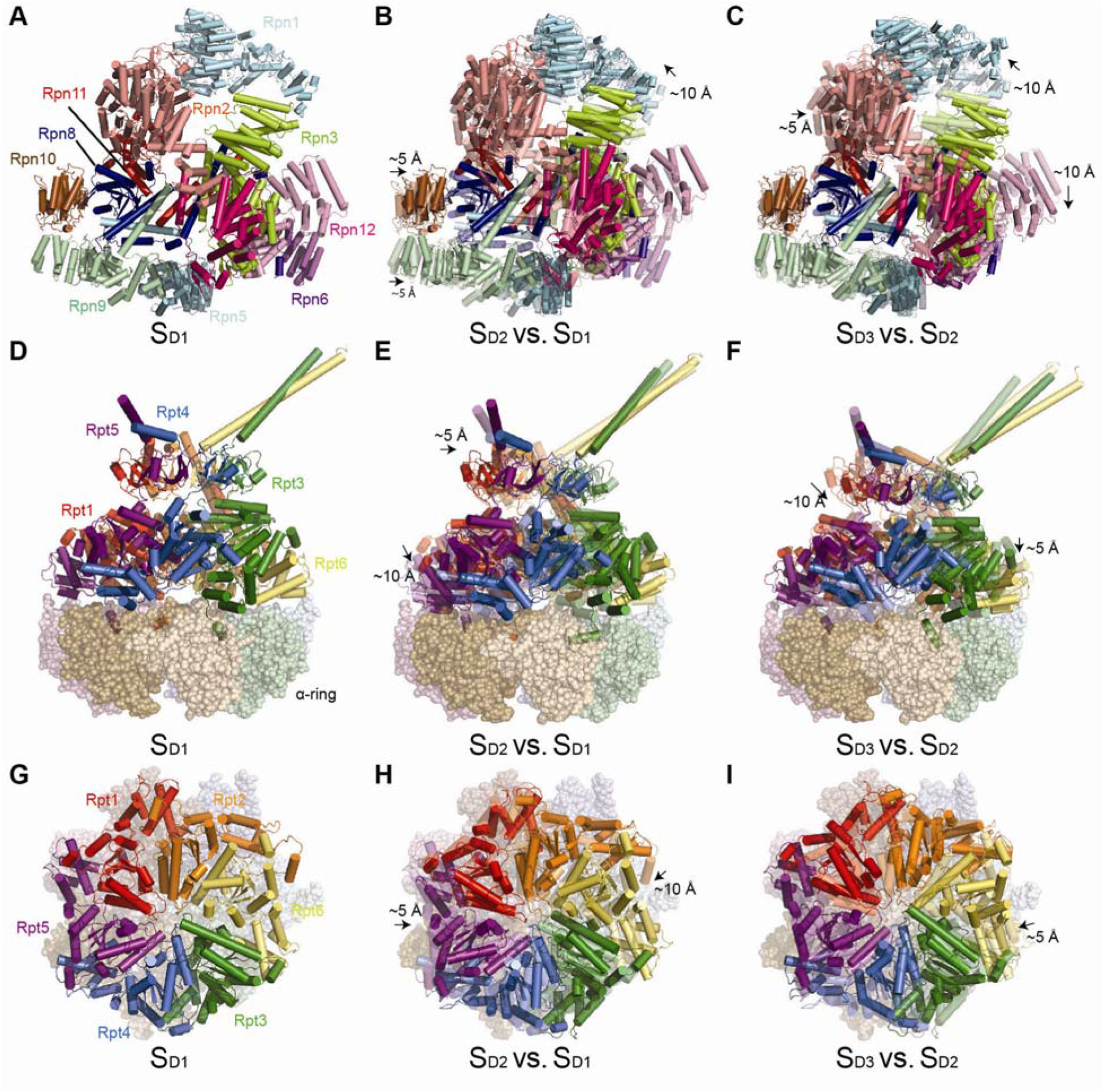
Conformational changes of the ATP-γS-bound human proteasome in three different S_D_-like states. (A) Top view of the lid of S_D1_. (B, C) Top view of the lid of S_D2_ (panel B) and S_D3_ (panel C) superimposed with the transparent cartoons of S_D1_ and S_D2_, respectively. (D) Side view of the AAA-ATPase ring above the α-ring in S_D1_. (E, F) Side view of the AAA-ATPase ring above the α-ring in S_D2_ (panel E) and S_D3_ (panel F) superimposed with the transparent cartoons of S_D1_ and S_D2_, respectively. (G) Top view of the AAA-ATPase ring in S_D1_. (H, I) Top view of the AAA-ATPase ring in S_D2_ (panel H) and S_D3_ (panel I) superimposed with the transparent cartoons of S_D1_ and S_D2_, respectively.

The lateral RP-CP interfaces, which are ancillary to regulate the conformations of the axial substrate-translocation pathway within the holoenzyme (Chen et al., 2016), also demonstrate prominent fluctuation. In the S_D1_ state, the N-terminal PCI domain of the lid subunit Rpn6 directly contacts the CP subunit α2 (Figure S5J). However, it is displaced ∼20 Å away from this interface in the S_D2_ state, leaving the Rpn6 PCI domain mostly dissociated from the α2-subunit (Figure S5K). This interface seems to be partly re-established in S_D3_ (Figure S5L); however, the N-terminal PCI domain of Rpn6 displayed a marked reduction in density as compared to that in S_D1_, suggesting instability of the interface in S_D3_ (Figure S5L).

### Rpn1 Engages with the AAA-ATPase Unfoldase through Coiled Coils

The Rpn1 subunit exhibits the lowest local resolution around 8-10 Å in the S_A_ map and so are the Rpn1 densities in all previously published high-resolution cryo-EM structures of proteasomes (Chen et al., 2016; Ding et al., 2017; Huang et al., 2016; Schweitzer et al., 2016; Wehmer et al., 2017). Notably, the local resolution of Rpn1 in the S_D1_ and S_D2_ maps are much improved over its counterpart in the S_A_ map, reaching 4.5-6.5 Å (Figure 4 and Figure S1). We thus retraced the Rpn1 backbone structure based on the new maps. After accounting all Rpn1 densities with a backbone model, there are extra densities displaying two neighboring helices located between Rpn1 and Rpt2 (Figure 4A,B). Given its proximity to the C-terminal of coiled coil domain of Rp1-Rpt2 dimer, we interpreted the extra densities as the N-terminal part of coiled coil of Rpt1-Rpt2 dimer.

**Figure 4.**
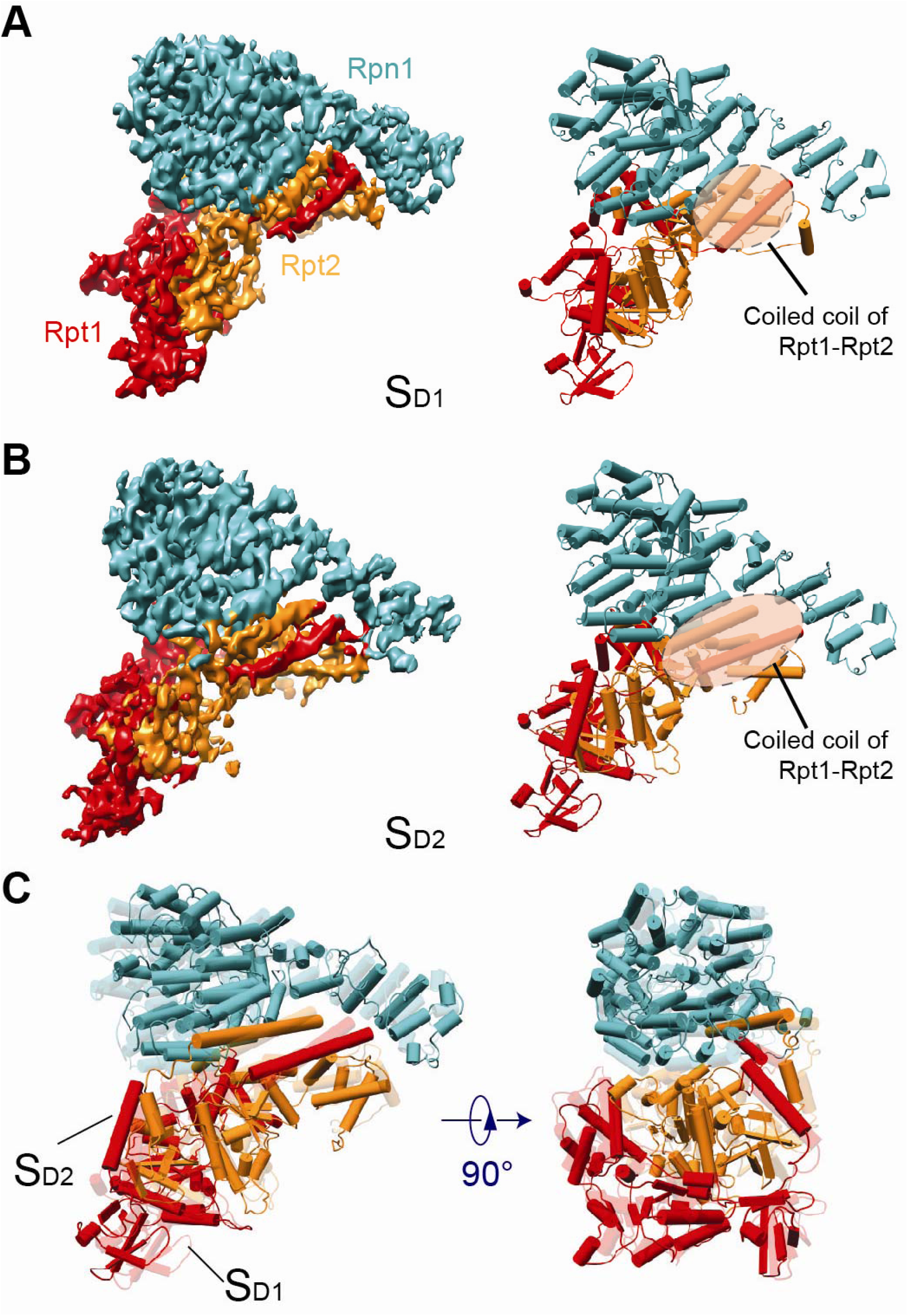
Rpn1 engages with the AAA-ATPase ring through the coiled coil. (A, B) The density maps (left) and atomic models (right) of the Rpn1 (cyan) interacting with Rpt1 (red) and Rpt2 (orange) in the AAA-ATPase ring in the S_D1_ and S_D2_ states, respectively, showing association between Rpn1 and the AAA-ATPase ring through coiled coils from Rpt1 and Rpt2. (C) Superimposition of the Rpn1-Rpt2-Rpt1 models in the S_D1_ and S_D2_ states in two different views.

The Rpn1 embraces a large interface with the Rpt1-Rpt2 coiled coil, with additional contacts with the AAA domain of Rpt2 stabilizing its association with the ATPase module (Figure 4A,B). Superposition of the Rpn1-Rpt2-Rpt1 structures in S_D1_ and S_D2_ indicates that the largest movement in Rpn1 lies in its N-terminal domain (Figure 4C). The movement of Rpn1 in the direction opposite to the motion of the AAA ring is conferred by the inter-domain rearrangement of the Rpt2 subunit, which is dependent of its nucleotide exchange in the state transition between S_D1_ and S_D2_ (see below).

### RP-CP Interface Stabilizes the Open Gate of the CP

In the published 8-Å map of the S_D_ state, we have observed that the C-terminal tails of Rpt subunits, except for Rpt4, are inserted into α-pockets (Chen et al., 2016). However, the resolution was too low to provide a reliable atomic modeling of the Rpt C-terminus in the previously reported S_D_ map. The counterpart state, s4, of the yeast proteasome had the similar resolution, leaving the perceived difference between the S_D_ and s4 states unexplained regarding the RP-CP interface (Wehmer et al., 2017). In the current study, the 3.5-Å map of the CP in the combined S_D_ state was substantially improved and thus allowed near-atomic model fitting of the C-terminal tail of Rpt subunits (Figure 5A-F). These C-terminal densities were also consistently observed in the cryo-EM maps of the S_D1_, S_D2_ and S_D3_ states at slightly lower resolutions. This observation suggests that the interactions between C-termini of Rpt subunits and α-pockets are invariant among S_D1_, S_D2_ and S_D3_. Our high-resolution structural data substantiate that the five C-terminal tails of Rpt1, Rpt2, Rpt3, Rpt5 and Rpt6 are inserted into the α-pockets to stabilize the open gate of CP.

**Figure 5.**
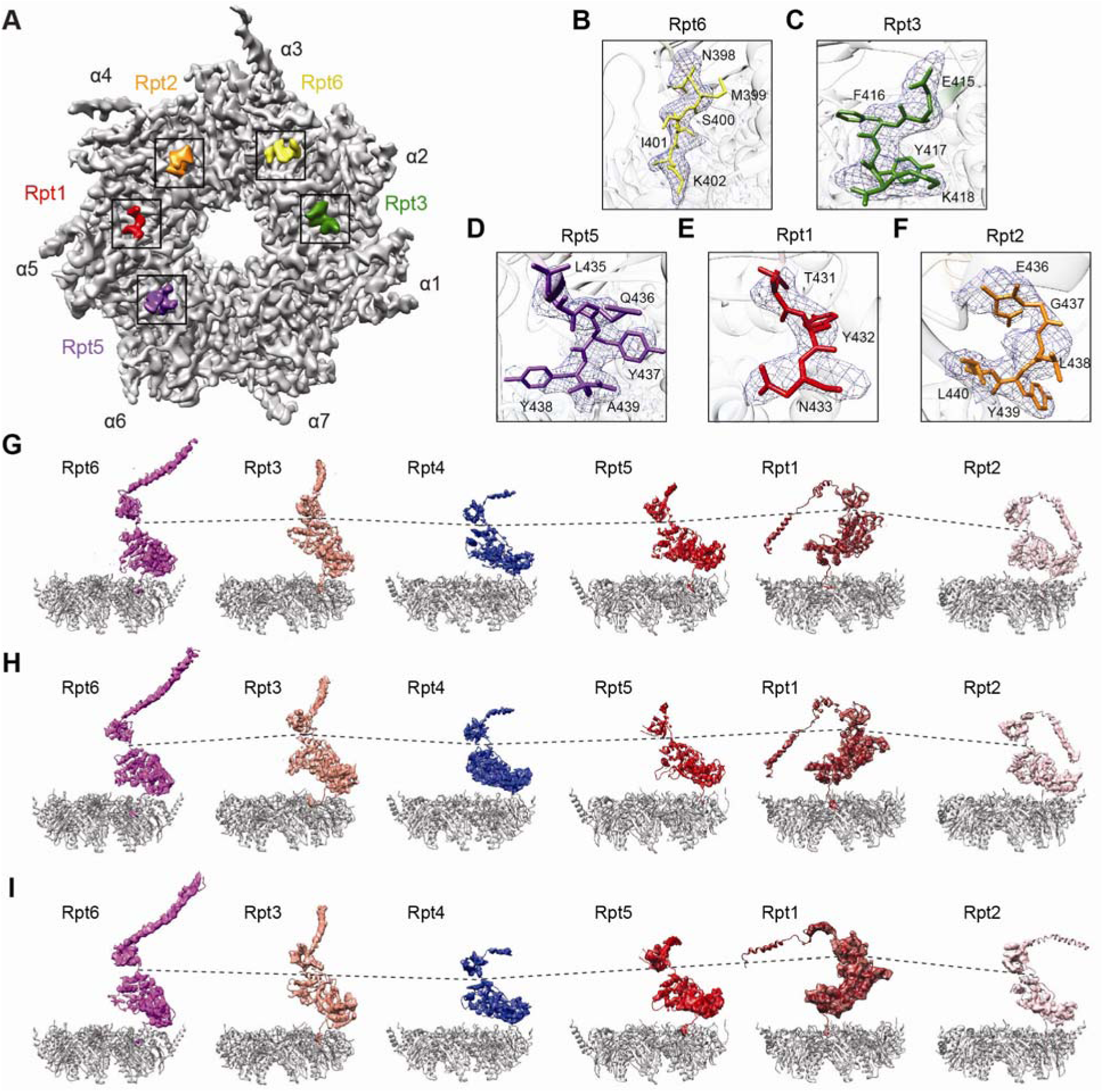
RP-CP interactions regulate the gate opening. (A) Overview of the RP-CP interface in which the C-terminal tails of Rpt1, Rpt2, Rpt6, Rpt3 and Rpt5 are shown to insert into the α-pockets in the S_D_ states. (B-F) Close-up views of each C-terminal tail shown in a stick representation superimposed with their corresponding cryo-EM densities shown in a mesh representation. (G-I) The Rpt subunits in a cartoon representation superimposed with cryo-EM densities showing the vertical positions of each Rpt subunit relative to the α-ring in a gray cartoon representation in the S_D1_ (panel G), S_D2_ (panel H) and S_D3_ (panel I) states.

### Structural Rearrangement in the AAA-ATPase Unfoldase

The AAA-ATPase ring in the resting state adopts a spiral “split lockwasher” architecture (Chen et al., 2016; Huang et al., 2016; Schweitzer et al., 2016), in which the spiral ring is split at Rpt6 with other Rpt subunits organized from the highest to the lowest position relative to the CP in the sequence of Rpt3-Rpt4-Rpt5-Rpt1-Rpt2. In the open-gate states, this spiral order with a single-split at Rpt6 is disarranged (Figure 5G-I). Specifically, Rpt1 raises to the height of Rpt3, while Rpt4 drops to the lower level around the position of Rpt2. Rpt6 also drops its level along with Rpt2 in S_D1_ and S_D2_, but raises to the higher level around the position of Rpt3 in S_D3_. Rpt2 demonstrates a prominent inter-domain motion between its small and large AAA subdomains, and falls back to the lowest position in S_D3_. The Rpt2 and Rp4 subunits compete for the lowest position, whereas Rpt1 and Rpt3 compete for the highest position. This rearrangement results in a novel AAA-ATPase ring architecture that follows a circle on a saddle topology. Along the circle, there is a two-period wave, with the highest Rpt3 and the lowest Rpt2 wobbling up and down as the ATPase ring vibrates laterally back and forth above the α-ring during conformational transitions among the three open-gate states.

### Nucleotide Exchange in the Open-Gate States

The nucleotide-binding pockets in the AAA family proteins share a common architecture surrounded by highly conserved motifs of Walker A, Walker B, sensor I, sensor II and arginine finger (Zhang and Wigley, 2008). The nucleotide binds the Walker A motif located next to a short linker between the small and large AAA subdomains of AAA-ATPases. The changes of nucleotide’s chemical state, among ATP, ADP-Phosphate, and ADP, modify the geometric relationship between the small and large AAA subdomain, thus regulating the conformations of the Rpt subunits (Ogura and Wilkinson, 2001; Sauer and Baker, 2011; Sledz et al., 2013). In both human and yeast proteasomes in the resting state, all six nucleotide binding sites are occupied (Chen et al., 2016; Ding et al., 2017; Huang et al., 2016; Schweitzer et al., 2016; Wehmer et al., 2017). Consistent with these observations, we identified nucleotide densities in all six Rpt subunits in the density map of the S_A_ state of our ATP-γS bound proteasome (Figure S6A,B).

In contrast to the full occupancy of nucleotide-binding sites in the S_A_ state, we found partial occupancy in the open-gate states. In the S_D1_ state, we can only observe nucleotide densities in Rpt1, Rpt3, Rpt4 and Rpt5. The absence of nucleotide densities in Rpt2 and Rpt6 suggests that these AAA-ATPase subunits are in an apo state with the nucleotides released (Figure 6A and Figure S6E). Similarly, in the S_D2_ state, we did not observe nucleotide densities in Rpt2 and Rpt3, but found them in Rpt1, Rpt4, Rpt5 and Rpt6 (Figure 6B and Figure S6F). The resolution of the S_D3_ state is not high enough to allow conclusive fitting of nucleotides. However, a tentative interpretation on the nucleotide-associated densities suggests that the nucleotide-binding sites in Rpt1, Rpt2, Rpt3 and Rpt5 are occupied (Figure S6C,D,G). The nucleotide-binding status in S_D3_ awaits further verification at a higher resolution. Thus, nucleotide release and exchange take place in Rpt2, Rpt3, Rpt6, and potentially Rpt4 as well, during conformational transitions among these open-gate states, and between the S_D_-like states and the gate-closed states such as S_A_.

**Figure 6.**
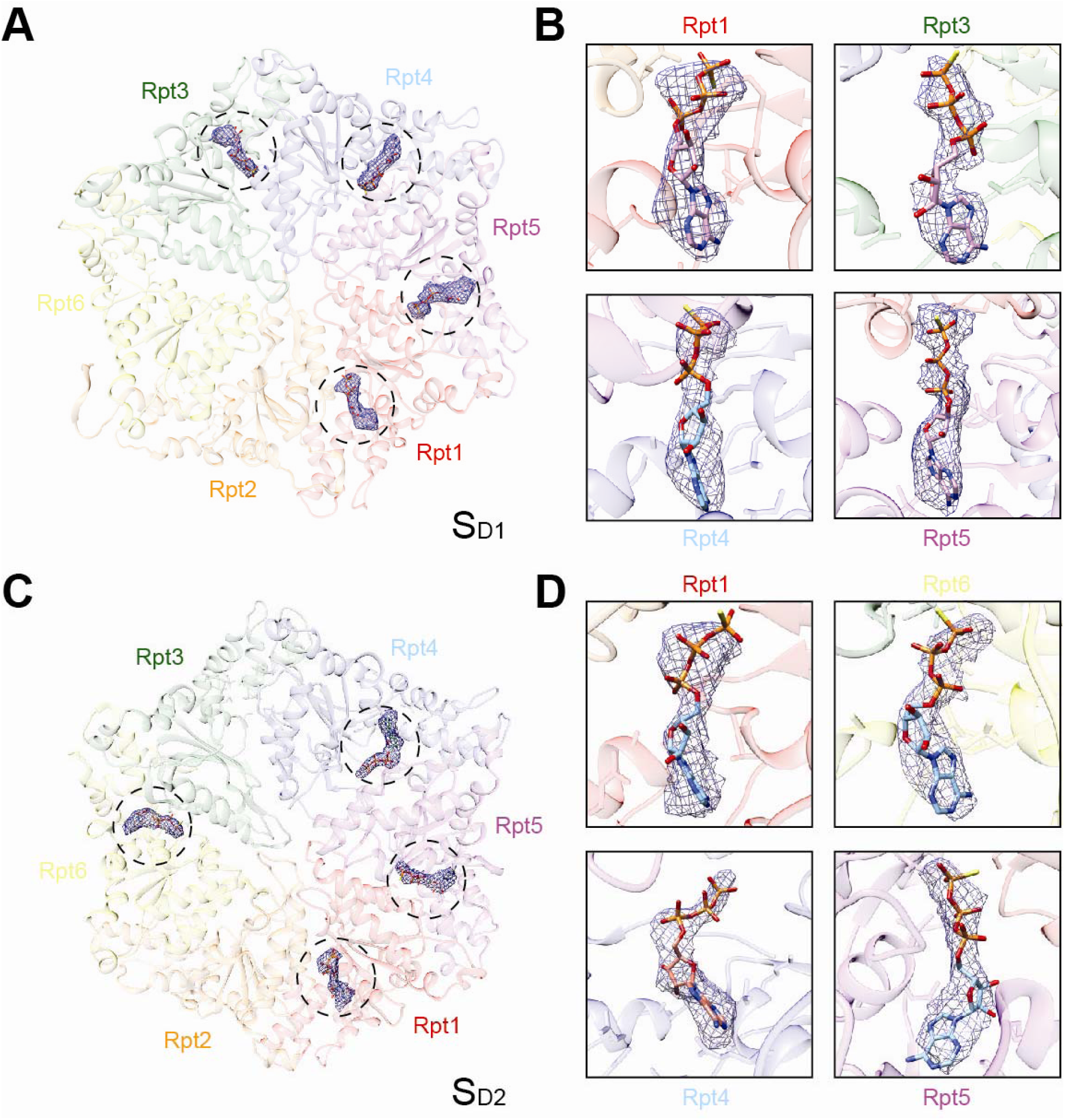
The cryo-EM densities of the nucleotides in the SD1 and SD2 states of the ATPγS-bound human 26S proteasome. (A, C) Overview of the nucleotide-binding sites in the AAA-ATPase heterohexamer of the ATP-γS-bound human 26S proteasome in the S_D1_ (panel A) and S_D2_ (panel C) states. Bound nucleotides are shown in a stick representation superimposed with their cryo-EM densities in a blue mesh representation. (B, D) Close-up views of nucleotide conformations in the nucleotide-binding sites in the S_D1_ (panel B) and S_D2_ (panel D) states. ATP-γS was modeled into the nucleotide densities of four Rpt subunits in each state.

The release of a nucleotide in an AAA-ATPase protein destabilizes the nucleotide-bound conformation, generating large structural arrangements between the small and large AAA subdomains (Glynn et al., 2009). Indeed, the Rpt2 and Rpt6 densities in the S_D1_ map, the Rpt2 density in the S_D2_ map, and the Rpt6 density in the S_D3_ map are of lower quality at slightly lower local resolutions than other Rpt subunits in the same map, indicating that their conformations are destabilized due to nucleotide release (Figure S6E-G).

### Topological Remodeling of the Substrate-Translocation Pathway

The central channel formed by the hexameric ring of the AAA domain of Rpt subunits is narrowed by inward-facing pore loops that were thought to drive the translocation of substrates (Beckwith et al., 2013; Glynn et al., 2009; Zhang et al., 2009a). In the S_A_ state, the pore-1 and pore-2 loops from neighboring subunits were found to pair with each other to constitute the constrictions of the AAA channel (Chen et al., 2016). The hydrophobic residues in the pore-1 loops of Rpt4, Rpt5 and Rpt1 are paired with charged residues in the pore-2 loops of Rpt3, Rpt4, and Rpt5, respectively (Chen et al., 2016). Although the AAA-ATPase in the S_D1_ state is in a conformation with its nucleotide-binding configuration quite different from that in S_A_, the pore-loop pairing architecture in S_D1_ resembles that in S_A_. By contrast, in the S_D2_ and S_D3_ states, the pore loops are dramatically reorganized into tilted, saddle-shaped circles (Figure 7). The saddle-like circle formed by the pore loops in S_D2_ is tilted toward the direction opposite to that in S_D3_, compatible with the overall wobbling motion of the AAA-ATPase ring.

**Figure 7.**
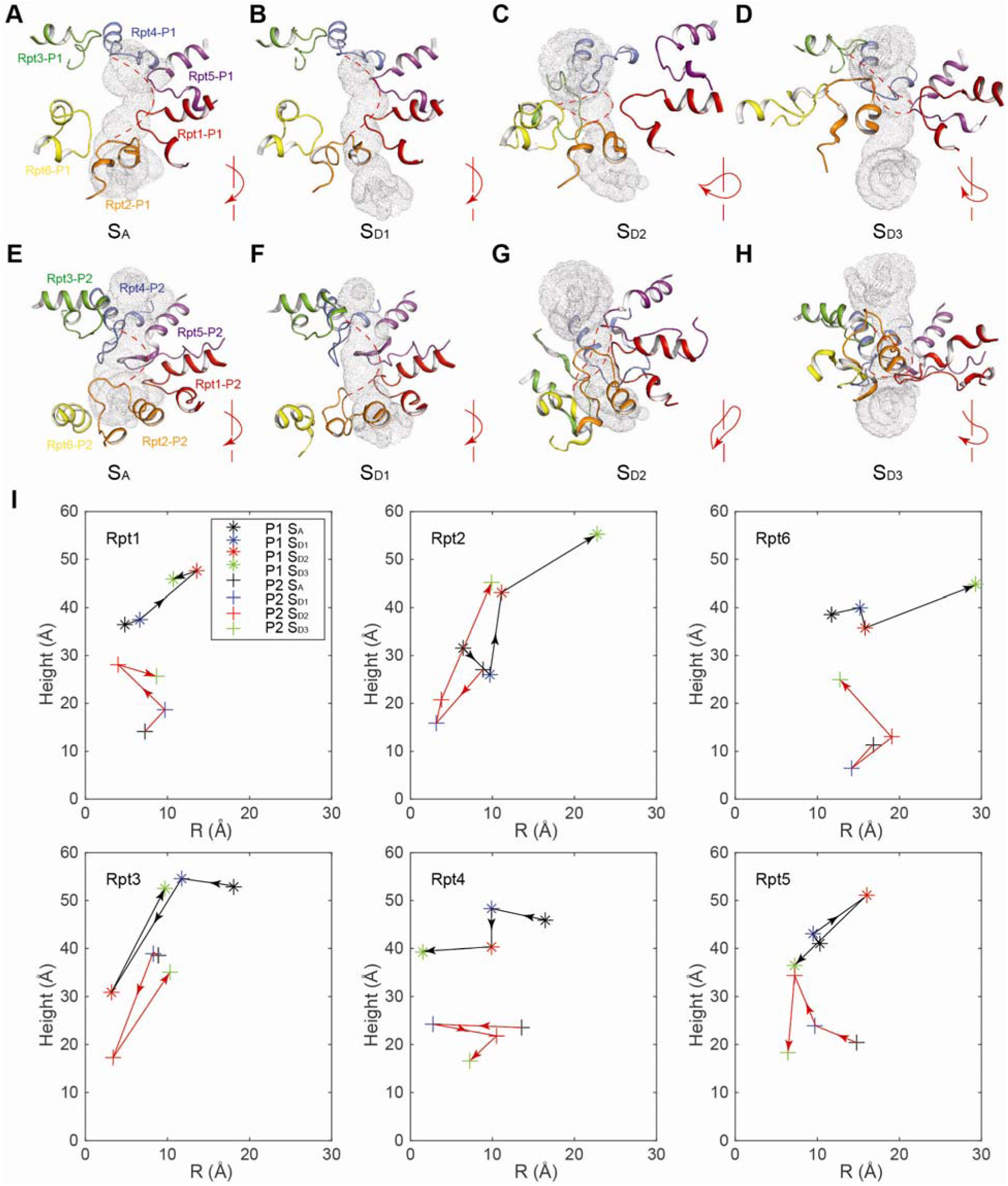
Topological remodeling of the pore loops along the substrate-translocation channel. (A) Close-up view of the pore-1 loops from six Rpt subunits decorating the AAA channel in S_A_ state. The solvent accessible surface of the AAA channel is estimated by program HOLE and is rendered with grey dots. The AAA channel is aligned top-down vertically, where substrates are supposed to enter from the top end of the pathway. (B-D) Close-up views of the pore-1 loops from six Rpt subunits decorating the AAA channel in S_D1_, S_D2_ and S_D3_ states. The lower right inset in each panel shows an illustrative graph summarizing the pore-1 loop organizing topology. The arrows show the clockwise direction of ATPase oligomeric organization from Rpt3, through Rpt4, Rpt5, Rpt1 and Rpt2, to Rpt6. The pore-1 loops in S_D1_ and S_D2_ show saddle-like topologies versus the spiral topology in S_D1_ and S_A_. The AAA channels are aligned to that shown in (A) based on their positions relative to the common reference of the CP. (E) Close-up view of the pore-2 loops from six Rpt subunits decorating the AAA channel in S_A_ state. (F-H) Close-up view of the pore-2 loops from six Rpt subunits decorating the AAA channel in S_D1_, S_D2_ and S_D3_ states. (I) The relative positional changes of each pore loop in S_A_, S_D1_, S_D2_ and S_D3_. The arrows show the direction of movements in the pore loops during state transitions from S_A_ to S_D1_, to S_D2_ and to S_D3_. Vertical axis indicates the relative height of the pore loops along the axis of the AAA-ATPase channel and the CP gate. Horizontal axis indicates the lateral distance orthogonal to the channel axis.

Driven by the nucleotide release in Rpt2 and Rpt3, the pore-1 and pore-2 loops in Rpt2 and Rpt3 are moved down in S_D2_ and then up in S_D3_, travelling over the longest distance of about 40-60 Å from S_D1_ to S_D3_ (Figure 7I). By contrast, both pore-1 and pore-2 loops in Rpt1 and Rpt4 travel down over much shorter distances (∼20-30 Å) in the state transition from S_D1_ to S_D2_ and to S_D3_. Most pore loops, more prominently in Rpt2, Rpt3 and Rpt5, alternate up and down in the triple-state transition. In contrast to vertical movements in most pore loops, the pore loops in Rpt4 mostly move laterally with distances around 10 Å.

In the S_A_ state, the pore-loop paring along the axial channel of the AAA ring makes a passage too narrow to smoothly translocate even an unfolded polypeptide (Chen et al., 2016; Huang et al., 2016; Schweitzer et al., 2016). The constrictions narrowed by the pore loops are substantially reconfigured in the S_D2_ and S_D3_ so that the AAA channel is overall widened relative to that in the S_D1_ state. However, the narrowest constrictions are still too narrow to allow translocation of peptides with large aromatic side chains such as tryptophan in S_D1_, S_D2_ and S_D3_ (Figure S7), suggesting that conformational flexibility in the pore loops might be necessary to accommodate substrate translocation (Glynn et al., 2009). Indeed, although the pore loops of Rpt6 do not contribute to the constriction of the AAA channel in S_D1_, they move in to directly shape the construction of the AAA channel in S_D2_. By contrast, the pore-1 loops of Rpt5 and Rpt6 are moved away from the substrate-translocation pathway in S_D2_ and S_D3_, respectively (Figure 7C, D), whereas the pore-2 loops of Rpt5 and Rpt6 remain engaged in the formation of the AAA channel constriction in S_D2_ and S_D3_ (Figure 7G, H). Hence, the architectural remodeling of the substrate-translocation channel observed in the three S_D_-like states manifests a marked conformational plasticity of the pore loops presumably necessary for substrate unfolding.

## DISCUSSION

In this work, we discovered three distinct open-gate states of the activated human proteasome at high resolution, expanding the number of known coexisting conformational states from four to six. Consistent with the previous study on the ATP-γS-bound yeast proteasome (Sledz et al., 2013), the conformational states of the human proteasome are redistributed upon the binding of slowly hydrolysable ATP-γS as opposed to ATP. All previous studies and the present work suggest that full occupancy of six nucleotide-binding sites stabilizes the resting state, where the CP gate is closed in both human and yeast proteasome holoenzymes (Chen et al., 2016; Ding et al., 2017; Huang et al., 2016; Schweitzer et al., 2016; Wehmer et al., 2017). Full occupancy of ATP-γS in all nucleotide-binding sites may be less favorable, allowing more copies of the complex to assume the S_D_-like states. Nonetheless, the S_A_ state is still the dominant population (∼50%) among the ATP-γS-bound human proteasome. For the first time, we observed the nucleotide exchange when the CP gate is open in an asymmetric fashion. We found that only four nucleotide-binding sites are occupied in the S_D_-like open-gate states. This is reminiscent of the previous studies suggesting that only four nucleotides are needed for the functioning of AAA-ATPase (Glynn et al., 2009; Horwitz et al., 2007). Similar to our S_D1_ state, a recent study on the yeast proteasome in complex with ADP-AlFx showed that the nucleotide release in Rpt2 and Rpt6 allows the AAA-ATPase to adopt a different conformation when the CP gate is still closed (Ding et al., 2017). Taken together, our results suggest that the release of at least two nucleotides may be necessary but insufficient for promoting the opening of the CP gate.

The high-resolution structures of the open-gate states clarify the mechanism of CP gate opening for the human proteasome. The open-gate s4 state of the yeast proteasome (Wehmer et al., 2017), resembling our previously reported S_D_ state (Chen et al., 2016), was recently visualized at relatively low resolution in the holoenzyme in complex with ADP-BeF_x_ and ATP/BeF_x_. In contrast to the human S_D_ state, the C-terminal tails of Rpt1 and Rpt4 were not seen to dock into the α-pockets in the yeast s4 state (Wehmer et al., 2017). This was at odds with our observation that only the Rpt4 C-tail is not inserted into the α-pockets in the open-gate human proteasome (Chen et al., 2016). Indeed, our high-resolution cryo-EM maps of all three open-gate S_D_-like states consistently visualized the densities of five Rpt C-tails in the α-pockets, confirming the C-tail insertion of all Rpt subunits except Rpt4 into the α-ring.

The multiple conformations of the AAA-ATPase unfoldase associated with an open CP gate provide novel mechanistic insights into the proteasome activation and initiation of substrate translocation. Engagement of substrates must be coordinated with distributed conformational changes in the RP and the RP-CP interface. Binding of six nucleotides in the AAA-ATPase ring may lock the topological relationship between the RP and the CP in the S_A_ state, and does not permit a continuous movement in the AAA-ATPase module. To allow more Rpt C-terminal tails to reach their nearby α-pockets, which seems to be crucial for CP gate opening, the structural constraints preventing the continuous movement in the base must be relieved. Thus, the human proteasome undergoes dramatic conformational changes in the RP throughout the S_B_ and S_C_ states to attain the S_D_-like states (Chen et al., 2016), in which the AAA-ATPase unfoldase gains certain freedom in simultaneously exploring different conformations that are compatible with the initiation of substrate translocation. In the S_D_-like states, the wobbling of the AAA ring is coincident with the lateral vibration of the whole AAA-ATPase ring, along with the vertical rocking of the OB ring. Given the similarity of pore-loop architecture between S_A_ and S_D1_, the S_D1_ state likely represents the conformation where an incoming substrate encounters the pore loops. Transition of S_D1_ to either S_D2_ or S_D3_ might promote the engagement of the AAA channel with an incoming substrate and the initiation of its translocation through vertical translation of the pore-loops in Rpt3. Thus, sustaining certain degree of AAA-ATPase dynamics in the absence of substrates, when the CP gate is open, might be necessary for axial engagement of substrates with the AAA channel and for a smooth transition into the substrate-translocation phase. We hypothesize that substrate binding with the pore-loops would further expand the conformational landscape of the AAA-ATPase unfoldase, driving it to a greater level of conformational dynamics to meet the functional needs of converting chemical energy of ATP hydrolysis into mechanical work on the axial, unidirectional movement of an engaged substrate. A full revelation of the functional cycle of the holoenzyme will require an in-depth analysis on the complete conformational spectrum and state sequence of the substrate-engaged proteasome at high resolution.

## EXPERIMENTAL PROCEDURES

### Protein Expression and Purification

Human proteasomes were affinity-purified on a large scale from a stable HEK293 cell line harboring HTBH (hexahistidine, TEV cleavage site, biotin, and hexahistidine) tagged hRPN11 (a gift from L. Huang, Departments of Physiology and Biophysics and of Developmental and Cell Biology, University of California, Irvine, California 92697) (Wang et al., 2007). The cells were Dounce-homogenized in a lysis buffer (50 mM NaH_2_PO_4_ [pH7.5], 100 mM NaCl, 10% glycerol, 5 mM MgCl_2_, 0.5% NP-40, 5mM ATP and 1mM DTT) containing protease inhibitors. Lysates were cleared, then incubated with NeutrAvidin agarose resin (Thermo Scientific) overnight at 4°C. The beads were then washed with excess lysis buffer followed by the wash buffer (50 mM Tris-HCl [pH7.5], 1 mM MgCl_2_ and 1 mM ATP). Usp14 was removed from proteasomes using wash buffer +150 mM NaCl for 30 min. 26S proteasomes were eluted from the beads by cleavage, using TEV protease (Invitrogen). The doubly-capped proteasome was enriched by gel filtration on a Superose 6 10/300 GL column at a flow rate of 0.15ml/minute in buffer (30 mM Hepes pH7.5, 60 mM NaCl, 1 mM MgCl_2_, 10% Glycerol, 0.5 mM DTT, 0.8 mM ATP). Gel-filtration fractions were concentrated to about 2 mg/ml. Right before cryo-EM sample preparation, the proteasome sample was buffer-exchanged into 50 mM Tris-HCl [pH7.5], 1 mM MgCl_2_, 1 mM ATP-γS, 0.5 mM TCEP, and was supplemented with 0.005% NP-40.

### Data Collection

CryoEM sample grids were prepared with FEI Vitrobot Mark IV. C-flat grids (R1/1 and R1.2/1.3, 400 Mesh, Protochips, CA, USA) were glow-discharged before a 2.5-μl drop of 1.5 mg/ml proteasome solution was applied to the grids in an environment-controlled chamber with 100% humidity and temperature fixed at 4 °C. After 2 sec blotting, the grid was plunged into liquid ethane and then transferred to liquid nitrogen. The cryo-grids were imaged in an FEI Tecnai Arctica microscope, equipped with an Autoloader and operating at an acceleration voltage of 200 kV at a nominal magnification of 235,000 times. Coma-free alignment was manually conducted prior to data collection. Cryo-EM data were collected semi-automatically by the Leginon (Suloway et al., 2005) version 3.1 on the Gatan K2 Summit direct detector camera (Gatan Inc., CA, USA) in a super-resolution counting mode, with 7.5 s of total exposure time and 250 ms per frame. This resulted in movies of 30 frames per exposure and an accumulated dose of 30 electrons/Å^2^. The calibrated physical pixel size and the super-resolution pixel size are 1.5 Å and 0.75 Å, respectively. The defocus in data collection was set in the range of −0.7 to −3.0 μm. A total of 10,369 movies were collected, among which 8,463 movies were selected for further data analysis after visual inspection of each image for quality.

### Cryo-EM Data Processing and Reconstruction

All frames of the raw movie were first corrected for their gain using a gain reference recorded within two days of the acquired movie, then shifted and summed to generate a single micrograph that was corrected for overall drift. Each drift-corrected micrograph was used for the determination of the actual defocus of the micrograph with the CTFFind3 program (Mindell and Grigorieff, 2003). Reference-free 2D classification were carried out in both RELION 1.3 and ROME 1.0 that combined the maximum-likelihood based image alignment and statistical machine-learning based classification (Wu et al., 2016). 3D classification and high-resolution refinement were conducted in RELION 1.3 (Scheres, 2012). 502,384 complete particles of proteasome picked using DeepEM, an in-house developed program based on a machine-learning algorithm (Zhu et al., 2016). The initial model was generated in EMAN2 (Tang et al., 2007). A subset of 15,044 particles was used for reference-free classification performed by e2refine2d.py and for the generation of an initial model using e2initialmodel.py.

All 2D and 3D classifications were done at the pixel size of 1.5 Å. After the first round of reference-free 2D classification, bad particle classes were rejected upon inspection of class average quality, which left 419,237 particles. The initial model, low-pass filtered to 60 Å, was used as the input reference to conduct unsupervised 3D classification into 4 classes without assumption of any symmetry, using an angular sampling of 7.5° and a regularization parameter T of 4. Three classes have both RP caps on while the other has a single RP cap. Two of the double-cap classes, accounted for 45% percent of the particles, showed an open-gate feature while the other double-cap class and the single-cap class appeared in a close state. The two groups were separated and each was further sorted by reference-free 2D classification, and a total of 395,941 good particles were selected from the two groups for the following analysis.

To reduce the irrelevant heterogeneity in our particle population due to conformational variations, and to improve the map resolution, each complete proteasome particle was split into two pseudo-single-cap particles by a mask including an RP and a complete CP. We re-centered each particle to its new center calculated from the refined Euler angles and x/y-shifts. Accordingly, the center of each particle was shifted toward its RP part. We then shrank the box size and re-extracted the particles from raw micrographs. There are 377,313 pseudo-single-cap particles in the dataset of the open-gate states and 461,161 pseudo-single-cap particles in the dataset of the gate-closed state. Both datasets were further sorted by reference-free 2D classification separately, and 370,068, 421,814 good particles left in the closed-and open-gate datasets, respectively. The datasets of open- and closed-gate states were then subjected to the second round of 3D classification separately, each into 6 classes. Five of the classes in the closed-gate states were similar in their overall conformations, which were combined with another two closed-gate classes classified from the “open-gate” dataset. The remaining one class from the closed-gate group has density that was partially averaged out, indicating large heterogeneity in the class. This class was further classified into four classes; three of which were abandoned and the other good class (29%) joined the two classes (40%) of the open-gate states from the second round the 3D classification. The new open-gate datasets subsequently went through another two rounds of 3D classification and gave rise to three S_D_-like states and a S_C_ state with 66,246, 75,726, 33,278 and 23,567 particles, respectively. Another round of 3D classification into four classes on the closed-gate dataset gave rise to the S_A_ state containing 241,329 particles in three classes. A collection of the remaining 68,819 particles from both groups was subjected to another round 3D classification into 4 classes and yields S_B_ states in the fourth class with 15,536 particles.

The final refinements of each state were done on the particle data at the counting mode with a pixel size of 1.5 Å. Based on the in-plane shift and Euler angle of each particle from the last iteration of refinement, we reconstructed the two half-maps of each state using single-particle images at the super-counting mode with a pixel size of 0.75 Å, which resulted in reconstructions with an overall resolution of 3.6 Å, 7.0 Å, 5.8 Å, 4.2 Å, 4.3 Å and 4.9 Å of the S_A,_ S_B,_ S_C,_ S_D1,_ S_D2_ and S_D3_ states, respectively, measured by gold-standard FSC at 0.143-cutoff on two separately refined half maps. Further, we combined S_D1,_ S_D2_ and S_D3_ datasets together and focused the refinement on the CP with a local mask, which yielded a 3.5-Å CP structure with one gate open and the other closed. We also applied focused refinement to improve the gold-standard resolution of the AAA-ATPase and CP component in the S_D1_, S_D2_ and S_D3_ states to 4.0 Å, 4.1 Å and 4.8 Å, respectively, by using a mask that kept the densities of AAA-ATPase and CP during the last several iterations of the refinement. Prior to visualization, all density maps were sharpened by applying a negative B-factor. Local resolution variations were further estimated using ResMap on the two half maps refined independently (Kucukelbir et al., 2014).

### Atomic Model Building and Refinement

To build the initial atomic model of S_A_ and S_D1-3_ states, we use previously published ATP-bound human 26S structure S_A_ and then manually improved the main-chain and side-chain fitting in Coot (Emsley and Cowtan, 2004) to generate the starting coordinate files. An exception is Rpn1 in S_D1-3_ states, where homology modeling was conducted in Modeller (Marti-Renom et al., 2000) using Rpn2 in the ATP-bound human 26S structure as a reference to generate a starting coordinate file. To fit the model to the reconstructed density map, we first conducted rigid-body fitting of the segments of the model in Chimera, and then the fit was improved manually in Coot. Finally, each atomic model refinement was carried out in real space with program Phenix.real_space_refine (Adams et al., 2010) with secondary-structure and geometry restrains to prevent overfitting. The pseudo-atomic models of S_B_ and S_C_ states were fitted in Coot starting from the ATP-bound human 26S structure S_B_ and S_C_, respectively, and were further refined in real space with Phenix.real_space_refine with secondary structure and geometry restrains.

### Structural Analysis and Visualization

Structural comparison was conducted in Pymol and Chimera. The solvent accessible surface of the peptide-conducting channels in different states were calculated by HOLE (Smart et al., 1996) with respect to the OB ring and the AAA ring separately. All figures of the structures were plotted in Chimera (Pettersen et al., 2004), Pymol (Harshbarger et al., 2015), or our Python code that employs the Matplotlib package.

## ACCESS NUMBERS

The single-particle reconstructions and atomic coordinates reported in this paper have been deposited in the Electron Microscopy Data Bank, www.emdatabank.org (accession nos. EMD-8662 for the CP structure of S_D_, EMD-8663, EMD-8664, EMD-8665, EMD-8666, EMD-8667, and EMD-8668 for S_D1_, S_D2_, S_D3_, S_A_, S_B_, and S_C_, respectively) and Protein Data Bank, www.wwwpdb.org (PDB ID codes 5VFO for the CP structure of S_D_, 5VFP, 5VFQ, 5VFR, 5VFS, 5VFT, 5VFU for the ATP-γS-bound holoenzymes of the S_D1_, S_D2_, S_D3_, S_A_, S_B_, and S_C_ states, respectively). The raw micrographs and particle data have been deposited in the Electron Microscopy Pilot Image Archive, www.ebi.ad.uk/emdb/ampiar (accession no. EMPIAR-10090).

## SUPPLEMENTAL INFORMATION

Supplemental information includes seven figures and one table and can be found with this article online.

## AUTHOR CONTRIBUTIONS

Y.M. and Y.L. designed this study. Y.L. purified and characterized the specimen. W.L.W. and Y.L. prepared the specimen for data collection. W.L.W. screened the specimen and collected cryo-electron microscopy data. W.L.W. and D.Y. pre-processed the micrograph data. Y.Z. performed initial reconstruction, processed the particle data, refined the density maps, and built the atomic models. Y.M. drafted and finalized the manuscript with inputs from Y.Z., W.L.W., Y.L., and Q.O. Y.Z. and Y.M. prepared all figures in the manuscript. Y.M. supervised this study. All authors contributed to data analysis and manuscript preparation.

## ACKNOWLEDGEMENTS

The authors thank Drs. Daniel Finley and Marc W. Kirschner for constructive discussions. This work was funded in part by a grant of the Thousand-Talent Plan of China (Y.M.), by an Intel academic grant (Y.M.), by the research funds at the Peking-Tsinghua Center for Life Science at Peking University (Y.M.), by a grant from National Natural Science Foundation of China 91530321 (Y.M., Q.O.). The cryo-EM experiments were performed in part at the Center for Nanoscale Systems at Harvard University, a member of the National Nanotechnology Coordinated Infrastructure Network (NNCI), which is supported by the National Science Foundation under NSF award no. 1541959. The cryo-EM facility was funded through the NIH grant AI100645, Center for HIV/AIDS Vaccine Immunology and Immunogen Design (CHAVI-ID). The data processing was performed in part in the Sullivan cluster at Dana-Farber Cancer Institute, which is funded in part by a gift from Mr. and Mrs. Daniel J. Sullivan, Jr., and in part with the high-performance computing platform at the Peking-Tsinghua Center for Life Science at Peking University.

